# Atmospheric deposition exposes pandas to toxic pollutants

**DOI:** 10.1101/079632

**Authors:** Yi-ping Chen, Ying-juan Zheng, Qiang Liu, Yi Song, Zhi-sheng An, Qing-yi Ma, Aaron M. Ellison

## Abstract

The giant panda (*Ailuropoda melanoleuca*) is one of the most endangered animals in the world, and it is recognized worldwide as a symbol for conservation. A previous study showed that wild and captive pandas were exposed to toxins in their diet of bamboo, but the ultimate origin of these toxins is unknown. Here we show that atmospheric deposition is the origin of heavy metals and persistent organic pollutants (POPs) in the diets of captive and wild Qinling giant pandas. Atmospheric deposition averaged 115 and 49 g⋅m^−2^⋅yr^−1^ at China’s Shaanxi Wild Animal Research Center (SWARC) and Foping National Nature Reserve (FNNR), respectively. Atmospheric deposition of heavy metals (As, Cd, Cr, Pb, Hg, Co, Cu, Zn, Mn and Ni) and POPs at SWARC was higher than at FNNR. Soil concentrations of the aforementioned heavy metals other than As and Zn also were significantly higher at SWARC than at FNNR. We conclude that efforts to conserve the Qinling subspecies of panda may be compromised by air pollution attendant to China’s economic development. Improvement of air quality and reductions of toxic emissions are urgently required to protect China’s iconic species.

## Introduction

The giant panda (*Ailuropoda melanoleuca* (David, 1869)) is one of the most endangered animals in the world and a worldwide symbol for conservation. Two strategies, developed in the last several decades, are now used to protect this flagship endangered species. One strategy uses *ex-situ* breeding in, for example, the zoos of Beijing and the seven breeding centers, established since the 1950s, of Wolong and Chengdu. The other strategy has been to establish natural conservation zones that preserve panda habitat: 50 conservation zones, with a total area > 20,000 km^2^, have been delimited (Zhang and Wei 2006). In these conservation zones, efforts are ongoing to reduce habitat destruction, logging, resource exploitation, and tourism, all of which threaten wild panda populations.

As China’s economy has developed rapidly, environmental problems have emerged.This trade-off of environmental quality for economic development was common in the developed nations (Seinfeld 2004), and in China has had predictable effects of particulate pollution influencing air quality, regional and global climates, and human health (Cao et al. 2012, Wang et al. 2014). For example, in 2013, China experienced extremely severe and persistent haze pollution: measurements of average daily concentrations of PM_2.5_ (particulate matter with an aerodynamic < 2.5-µm diameter) in 74 major cities exceeded the Chinese pollution standard of 75 µg/m^3^. During the same year, a maximum daily concentration of 772 µg/m^3^ was observed over 1.3 million km^2^, affecting at the health of at least 800 million people (China National Environmental Monitoring Centre 2013).

Xi’an, one of the largest cities in China, is situated on the Guanzhong Plain at the Northern edge of the Qinling Mountains. This city has a resident population of eight million people and receives at least two million visitors annually. Between 2005 and 2010, the 24-hr PM_2.5_ in Xi’an ranged from 130-351 µg /m^3^ (Han et al. 2010, Shen et al. 2011), exceeding Chinese government standards 2-5-fold. Intense “haze-fog” events occur regularly, making air pollution one of the most important environmental issues in Xi’an (Cao et al. 2011).

Xi’an also is home to the Shaanxi Wild Animal Research Center (SWARC: 34° 06′N, 108° 32′ E). Established in 1987, SWARC is on the north slope of the Qinling Mountains and is dedicated to the conservation of the golden monkey (*Rhinopithecus roxellana* Milne-Edwards, 1870), golden takin (*Budorcas taxicolor* Hodgson, 1850), crested ibis (*Nipponia nippon* (Temminck, 1835)), and the Qinling subspecies of giant panda, of which only 345 individuals remain (Sun et al. 2005, SFA 2015). Captive pandas at SWARC and wild pandas elswhere in the region are exposed to heavy metals and persistent organic pollutants (POPs), including PCBs (polychlorinated biphenyls), PCDDs (polychlorinated dibenzo-p-dioxins), and PCDFs (polychlorinated dibenzofurans) through their diet of bamboo (Chen et al. *in press*). However, the ultimate origin of these pollutants is not known. Here we test the hypothesis that these pollutants are derived from atmospheric deposition.

## Methods and materials

### Sample collection

Atmospheric deposition samples were collected from November 8, 2013 to November 8, 2014 at the Foping National Nature Reserve (FNNR: Qinling Mountain, 33° 33′ – 33° 46′ N, 107° 40′ – 107° 55′E), Shaanxi Wild Animal Research Center (SWARC: Louguantai, Zhouzhi County, Xi’an city, 34° 06′ N, 108° 32′ E), and Xi’an City (34° 23′ N, 108° 89′ E). Samples of dry deposition and precipitation were collected continuously for one year into 66×40×12-cm plastic containers located at four sites at FNNR, three at SWARC, and four in Xi’an city. During the sampling period, purified water was added to the containers to avoid the collected deposition being blown out of the containers. After collection, the containers were rinsed with purified water to release particles deposited or sorbed onto the container walls. At the same time, soils samples also were collected from FNNR and at SWARC where bamboos are planted to feed captive pandas. Both the suspensions and the soil samples were dried to a constant weight at 60 ºC before being homogenized with a ball mill.

### Heavy metal analysis

Five hundred mg of each sample was placed into a Teflon digestion vessel to which was added 11 mL GR-grade acid digestion mixture (1mL HNO3, 3mL HCl, 5mL HF, 2mL HClO_4_) for digestion with an electric hot plate. After digestion, samples were diluted to 50 mL with ultrapure water (18.2 MΩ/cm^2^ Milli-Q water, Millipore). Heavy metal concentrations were analyzed using atomic absorption spectroscopy (AAS, ZEEnit 700P, Analytik Jena, Germany). Concentrations of Cu, Zn, Mn and Cr were measured using the air-acetylene flame method with electrically modulated deuterium-HCl background correction. The hydride-forming elements As and Hg were measured using the HS55 Hydride System. Concentrations of Cd, Ni and Pb were measured using a graphite furnace AAS coupled to a MPE 60 graphite autosampler with 2-field mode Zeeman effect background correction. Heavy metal concentrations are expressed as μg/g^1^ dry weight.

### Analysis of persistent organic pollutants

Sample extraction, cleanup, and chemical analysis of POPs followed established methods with some modifications (Liu et al. 2006, Chen et al. 2016, Li et al. 2008). Samples from atmospheric deposition were freeze-dried before being spiked with ^13^C-labeled surrogate standards (Environmental Protection Agency (EPA) methods 1613B and 1668A) and undergoing accelerated solvent extraction with dichloromethane: hexane (1:1). Each sample extract was adjusted to 50 ml with hexane; 15 g of acid silica (30% w/w) was added to remove lipids. The acid silica was stirred for 2 h and the extract was poured through ≈5 g of anhydrous sodium sulfate. All extracts were concentrated to 2 ml by rotary evaporation before cleanup. All solvents were purchased from Fisher (Fair Lawn, New Jersey, USA). Silica gel was obtained from Merck (silica gel 60, Darmstadt, Germany). Basic alumina was obtained from Aldrich (Brockmann I, standard grade, Milwaukee, Wisconsin, USA). Florisil was obtained from Riedel-de Haën (60–100 mesh ASTM, Seelze, Germany).

PCBs, PCDDs, and PCDFs were analyzed by the POP laboratory of the Research Center for Eco-environmental Sciences, Chinese Academy of Sciences; all concentrations were corrected for lipid weight. Twenty-five PCB congeners, including 12 dioxin-like congeners, were quantified with an isotope dilution method using high-resolution gas chromatography coupled with high-resolution mass spectrometry (HRGC/HRMS). Total organic carbon (TOC) concentration was analyzed on a TOC Analyzer (O.I Analyzer, College Station, Texas, USA). A 0.1 g sample was weighed and loaded into the combustion cup, which was packed with quartz wool. Prior to combustion, the samples were wetted with 5% phosphoric acid and heated to 250°C for 1 min to purge inorganic carbon. The signal was detected by non-dispersed infrared (NDIR) detection when flashed at 900 °C for 6 min in the combustion chamber. Calibration standard solutions, ^13^C_12_-labeled surrogate standards, and ^13^C_12_-labeled injection standards were purchased from Wellington Laboratories (Guelph, Canada).

Quantification of 17 PCDD and PCDF homologues was done by HRGC/HRMS on an Agilent 6890 gas chromatograph coupled with an Autospec Ultima mass spectrometer (Waters Micromass, Manchester, UK) operating in EI mode at 35 eV; the trap current was 600 Å. The GC was equipped with a CTC PAL autosampler. One or two microlitre samples were injected in splitless mode (splitless time, 2 min for PCDD/Fs) in a DB-5MS fused silica capillary column (60 m for PCDD/Fs and PCBs) with helium as carrier gas at a constant flow rate of 1.2 ml/min. The oven temperature programs were as follows: for PCDD/Fs, start 150 °C held for 3 min, 150-230 °C at 20 °C min^−1^ held for 18 min, 230-235 °C at 5 °C min^−1^ held for 10 min, 235-320 °C at 4°C min^−1^ held for 3 min; for PCBs, start 120 °C held for 1 min, 120-150 °C at 30 °C min^−1^, 150-300 °C at 2.5°C min^−1^ held for 1 min.

### Statistical Analysis

All statistical analyses were done using the SPSS 20.0 software (IBM SPSS Statistics, IBM Corp.,USA Inc.); the significance level was set at *P* < α = 0.05. Amounts of atmospheric heavy metals deposition from FNNR, SWARC, and Xi’an City were compared using one-way ANOVA followed by Tukey post-hoc tests. Comparisons of heavy metals concentrations in soils were done using *t*-tests. Because PCCD, PCDF, and PCB congeners differ in toxicity, toxic equivalency factors (set by the World Health Organization), were used to calculate a single toxic equivalent (WHO-TEQ) for each sample (Van et al. 2006).

## Results

The annual average rate (2013-2014) of atmospheric deposition of dust was 199 ± 6.50 g⋅m^−2^⋅yr^−1^ in Xi’an city, 115 ± 9.84 g⋅m^−2^⋅yr^−1^ at SWARC, but only 49 ± 6.79 g⋅m^−2^⋅yr^−1^ at FNNR. Deposition rates of all assayed heavy metals were significantly lower at FNNR than at SWARC (Fig. 1), and all but As were significant lower at SWARC than at Xi’an. In parallel, concentrations of all assayed heavy metals except for As and Zn in soils around SWARC were significantly higher than in soils around FNNR (Fig. 2). Concentrations of Cd, Pb, Zn and Mn at both SWARC and FNNR exceeded their soil background criteria (CNEMC 2000), whereas concentrations of Hg exceeded its soil background criterion only at SWARC (Fig. 2). There were significant positive correlations in the concentrations of these metals in deposited dust between Xi’an and SWARC (*r* = 0.98), and between Xi’an and FNNR (*r* =0.87).

**Figure 1.**
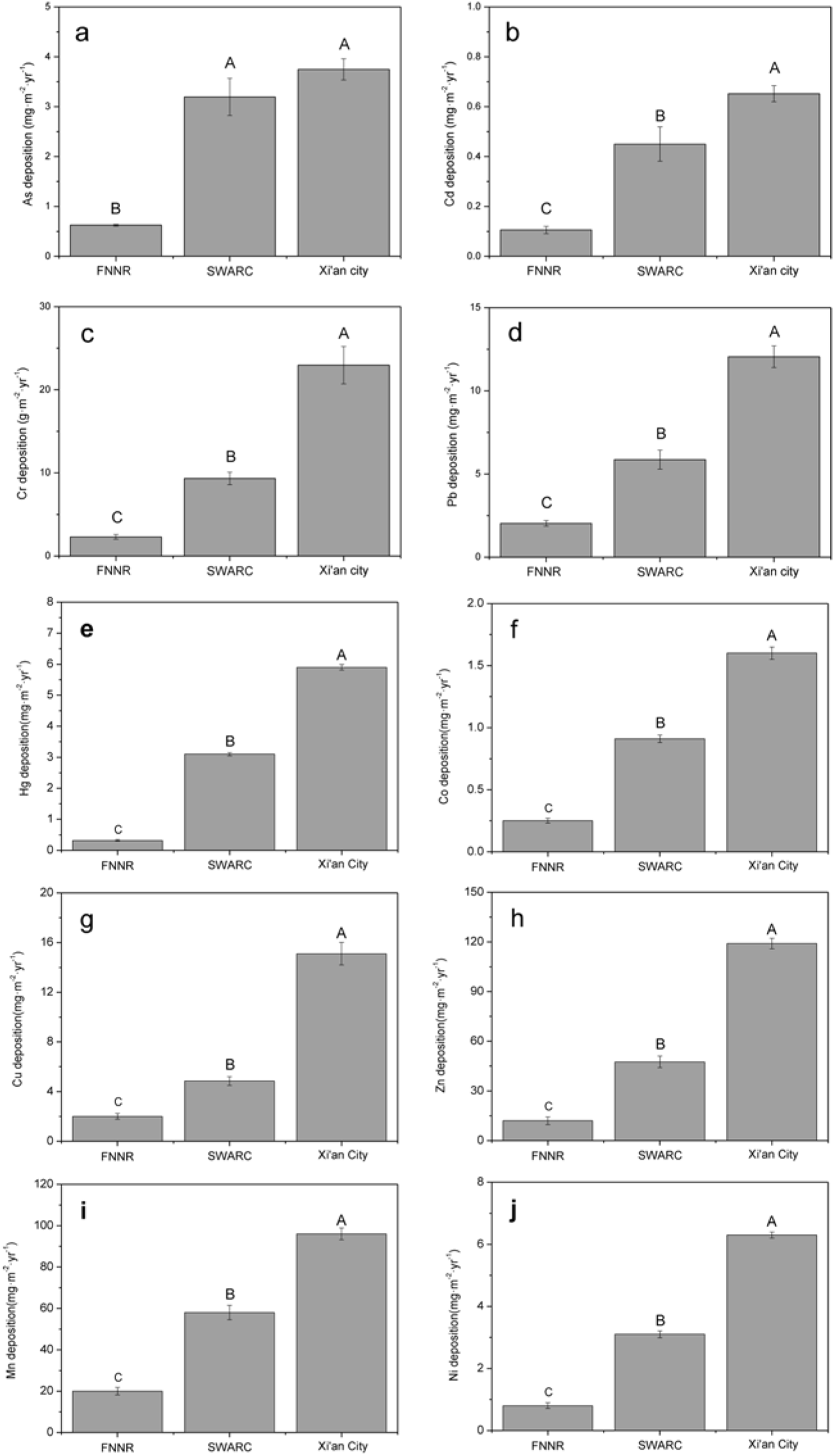
Mean (± 1 SE of the mean) amounts of atmospheric deposition of heavy metals: As (a), Cd (b), Cr (c), Pb (d), Hg (e), Co (f), Cu (g), Zn (h), Mn (i) and Ni (j) at the three studied sites [FNNR (n=4), SWARC (n=3) and Xi’an city (n=4)] over a one-year period. Differences among means at the three sites were compared using one-way AVOVA; different letters denote significant differences (*P* < 0.05; Tukey post-hoc test).

**Figure 2.**
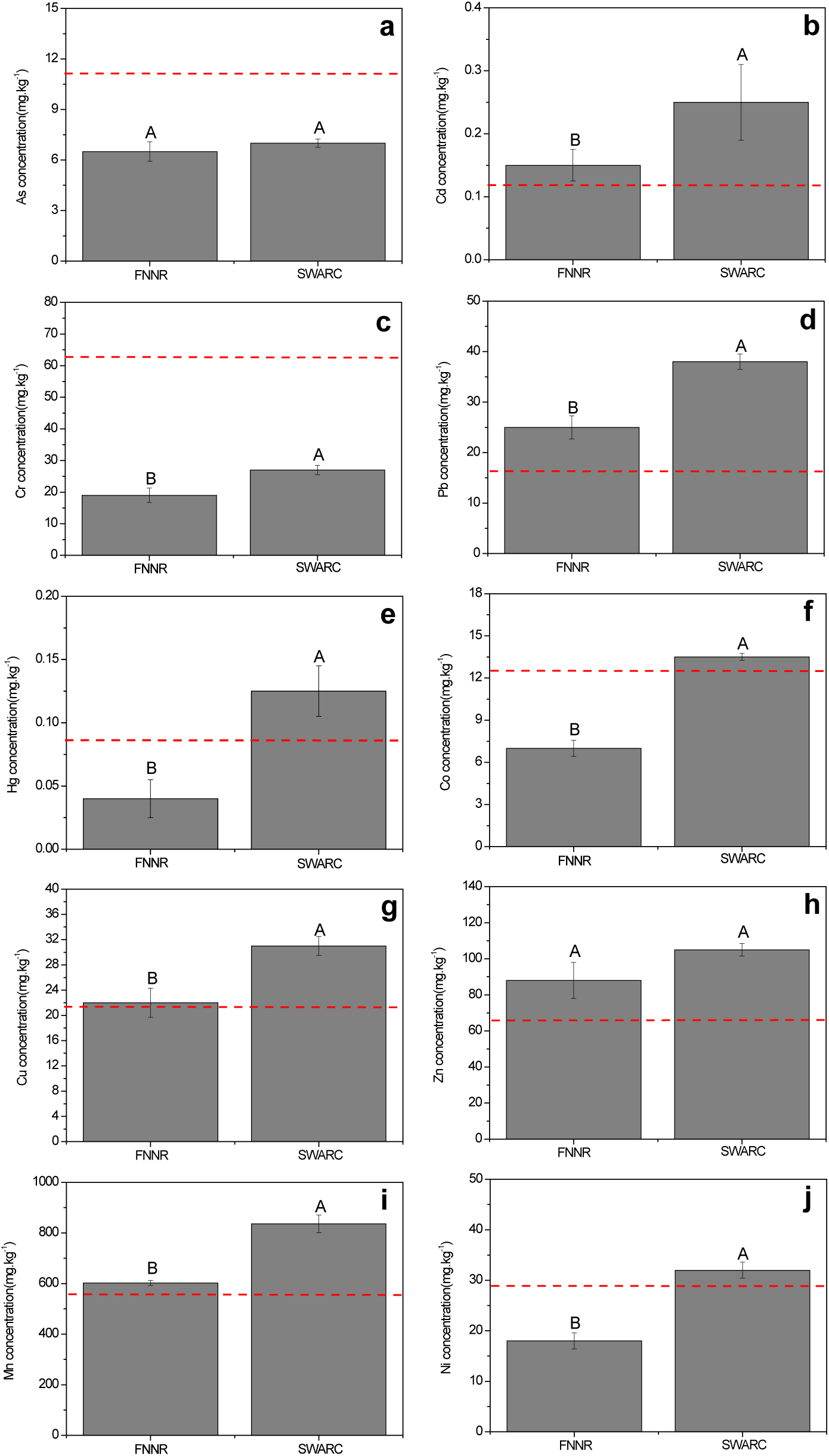
Comparison of toxic metals concentrations in soils at the southern (FNNR) and northern (SWARC) slopes of the Qinling Mountains. Bars illustrate mean (± 1 SE of the mean) amounts of soil As (a), Cd (b), Cr (c), Pb (d), Hg (e), Co (f), Cu (g), Zn (h), Mn (i) and Ni (j) at FNNR (n=4) and SWARC (n=3)]. A dashed red line in each panel indicates the soil background criteria value for polluted soils (CNEMC, 1990). Different letters denote significant differences (*P* < 0.05; *t*-test).

Deposition rates of dioxin and dioxin-like compounds (PCDDs and PCDFs) and PCBs were highest in Xi’an, intermediate at SWARC, and lowest at FNNR (Fig. 3). Seventeen congeners of PCDD/Fs and 12 of PCBs were detected in atmospheric deposition of PM (Table 1). The most prevalent PCDD/Fs were 1,2,3,4,6,7,8-HeptoCDF, OctaCDF and 1,2,3,4,6,7,8-HeptaCDD, whereas the most prevalent PCBs were 3,3',4,4'-TetraCB,2,3,3'4,4'-PentaCB, 2,3',4,4',5-PentaCB. The WHO-TEQ for PCDD/Fs and PCBs (Fig. 3c, 3d) paralleled trends in atmospheric deposition rates of total PCDD/Fs and PCBs (Figs. 3a, 3b).

**Figure 3.**
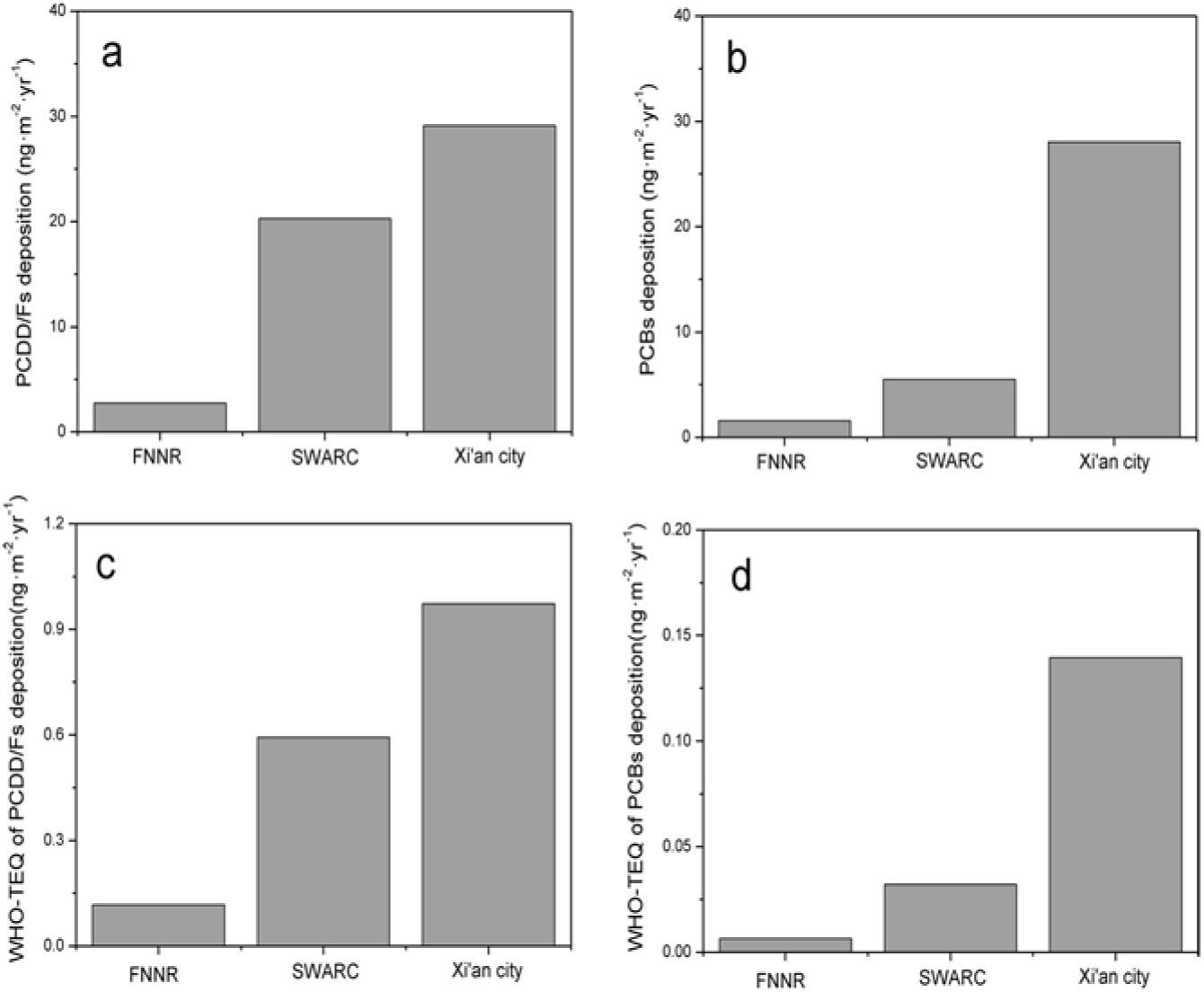
Total atmospheric deposition of PCDD/Fs (a) and PCBs (b), and the WHO-TEQs of PCDD/Fs (c) and PCBs (d) at the three different sites (FNNR, SWARC and Xi’an city) over a one-year period. Values are from four pooled samples for FNNR and Xi’an city, and three pooled samples for SWARC. Data for individual congeners of PCBs, PCDDs, and PCDFs are given in Table 1.

**Table 1.**
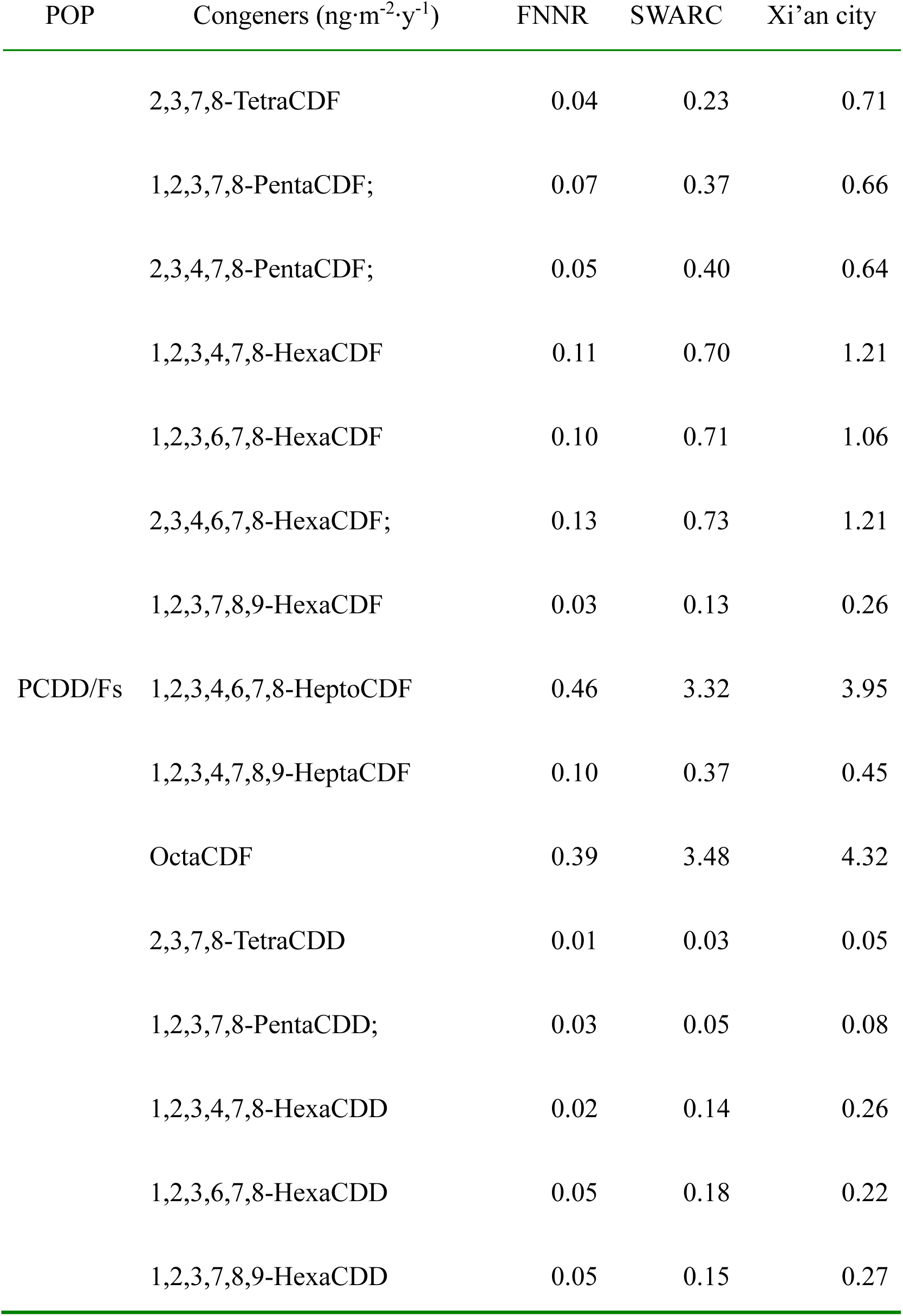
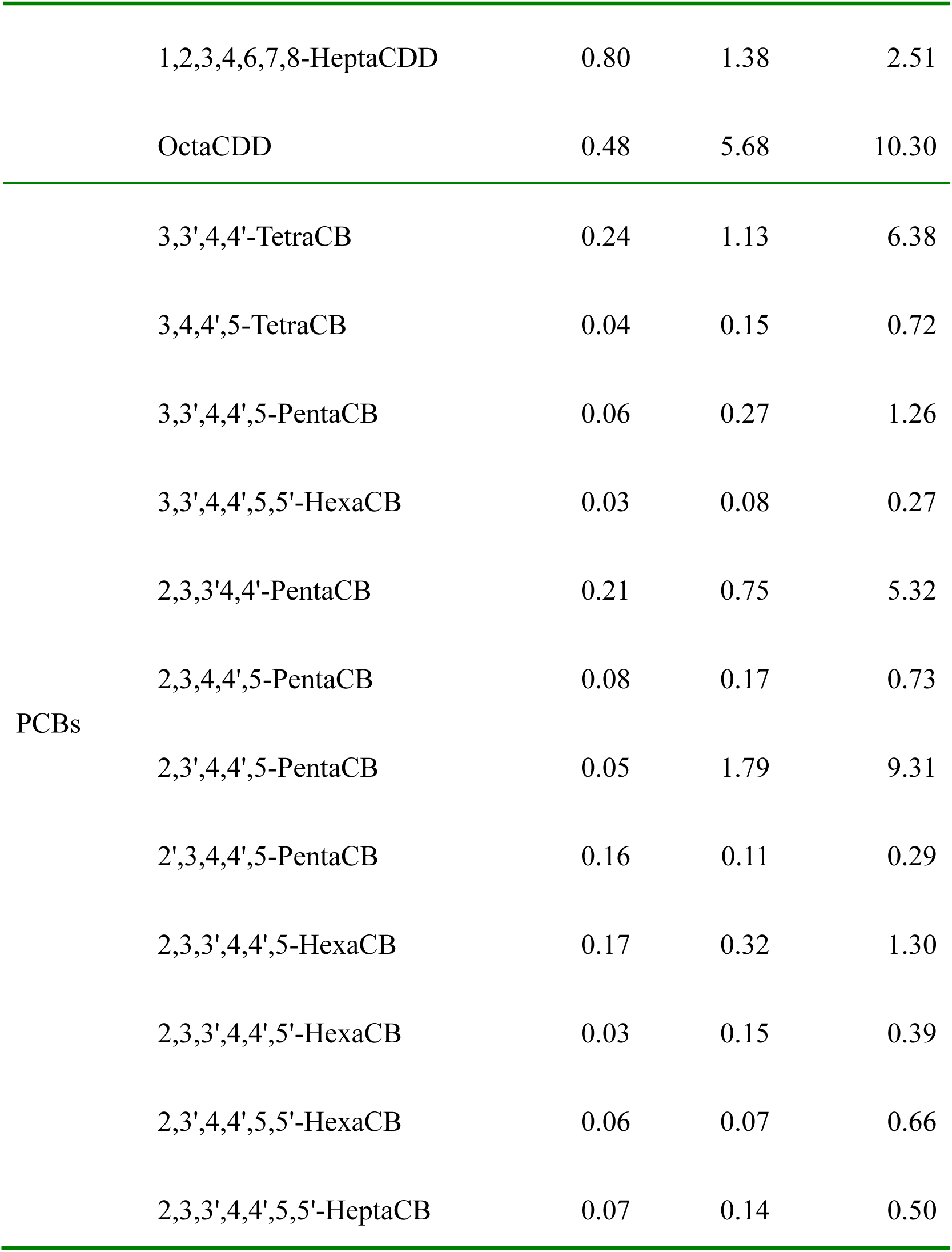
Concentrations of PCDD/F and PCB congeners in atmospheric deposition at the 295 different sites (FNNR, SWARC and Xi’an City). Data are from four pooled samples 296 from FNNR and Xi’an city, and three pooled samples from SWARC.

## Discussion

The Qinling Mountain region is home to a number of threatened and endangered species, including the golden monkey, golden takin, crested ibis, and the Qinling subspecies of the giant panda. The Shaanxi Wild Animal Research Center, which focuses on *ex situ* conservation of these species, is on the north slope of the Qinling Mountains and near Xi’an city. Intense “haze-fog” events have taken place in this region many times in recent years, and air pollution has become one of the important environmental issues in Xi’an. We explored possible relationships between atmospheric deposition of heavy metals and POPs in light of the previously documented exposure to all of these toxins in the diet of pandas (Chen et al. *in press*) and to some heavy metals in captive monkeys at SWARC (Liu et al. 2015).

The high annual deposition rate of metal- and pollutant-laden dust at Xi’an and SWARC originates from coal combustion; the transport of sand from deserts during droughts and from bare soil surfaces in the surrounding areas; heavy traffic; and a large number of on-going construction sites in the cities. The elevated levels of metals in deposition parallels that found in bamboo fed to captive pandas at SWARC (Chen et al. *in press*) and far exceeds that found in both deposition (Fig. 1–2) at and bamboo eaten by pandas in the wild (at FNNR) (Chen et al. *in press*). Correlations in the concentrations of metals between Xi’an city and either SWARC or FNNR suggest a source for these metals in the industrial activity and traffic in Xi’an (Dun and Tan 2013, Ha et al. 2014), upwind from both SWARC and FNNR. At high concentrations, these metals all have serious health effects (e.g., Rodier 1955, Friberg et al. 1985, Buchet and Lauwerys 1989, Winge and Mehra 1990, Rowbotham et al. 2000, Falcón et al. 2003, Doreswamy et al. 2004), and ten serious events of heavy metal (Hg, Cr, Cd, Pb and As) contamination in China have taken place in the past decade with significant public health impacts (Lu et al. 2015).

In parallel, deposition rates of dioxin and dioxin-like compounds (PCDDs and PCDFs) and polychlorinated biphenyls (PCBs) were highest in Xi’an, intermediate at SWARC, and lowest at FNNR (Fig. 3). Like heavy metals, PCDDs and PCDFs are by-products of combustion and industrial processes (Fiedler et al. 2007); these persistent organic pollutants are known human carcinogens and endocrine disruptors (Mai et al. 2005, van den Berg et al. 2006, Imamura et al. 2007). In contrast, PCBs were once used widely as non-flammable insulators and heat-exchange fluids (De Voogt et al. 1990), but their production ceased in 1974. Nonetheless, their long-term and continuing persistence in the environment and in tissues of living organisms has been associated with reduction in reproductive success, birth defects, and behavioral changes (Mai et al. 2005, Imamura et al. 2007).

The most prevalent congeners of PCDD/Fs recovered in samples of atmospheric deposition were 1,2,3,4,6,7,8-HeptoCDF, OctaCDF and 1,2,3,4,6,7,8-HeptaCDD, whereas the most prevalent PCBs were 3,3',4,4'-TetraCB, 2,3,3'4,4'-PentaCB, 2,3',4,4',5-PentaCB (Table 1). Total Concentrations of PCDD/Fs were higher at Xi’an than SWARC. Together, these data indicated that the main toxic organic chemicals deposition at SWARC originated from Xi’an.

Atmospheric deposition of pollutants in China, including in and around Xi’an, has been increasing rapidly (Cao et al. 2011). Deposition of heavy metals and POPs has resulted in high concentrations of these toxins in soils, well in excess of established soil background criteria. POPs and heavy metals can be inhaled directly by pandas or taken up by bamboo and subsequently bioaccumulate in pandas. Our results suggest that urban and industrial areas are the main sources of these environmental toxins, and pandas in captive breeding centers near cities are at greater risk than pandas in natural reserves further from urban areas and industrial centers. Rapid action to improve atmospheric conditions, including efforts to decrease automobile emissions, reduce coal usage, and improve urban efficiency should parallel efforts to relocate pandas from urban-based captive breeding centers to environmentally cleaner areas.

## Acknowledgments

This work was supported by funds from the IEECAS.

## Author’s contributions

Yi-ping Chen conceived the study and wrote the initial draft. Qiang Liu and Ying-juan Zheng performed the experiments, Yi Song performed statistical analysis, Zhi-sheng An contributed significantly to discussion, Qing-yi Ma performed sample collection, Aaron M. Ellison discussed results and their interpretation, and edited the manuscript.

